# Using 3D geometric morphometrics to aid taxonomic and ecological understanding of a recent speciation event within a small Australian marsupial (genus *Antechinus*)

**DOI:** 10.1101/2021.04.28.441717

**Authors:** Pietro Viacava, Andrew M. Baker, Simone P. Blomberg, Matthew J. Phillips, Vera Weisbecker

## Abstract

Taxonomic distinction of species forms the foundation of biodiversity assessments and conservation priorities. However, traditional morphological and/or genetics-based taxonomic assessments frequently miss the opportunity of elaborating on the ecological and functional context of species diversification. Here, we used 3D geometric morphometrics of the cranium to improve taxonomic differentiation and add eco-morphological characterisation of a young cryptic divergence within the marsupial carnivorous genus *Antechinus*. Specifically, we used 168 museum specimens to characterise the recently proposed clades *A. stuartii* “south”, *A. stuartii* “north” and *A. subtropicus*. Beyond slight differences attributable to overall size (and therefore not necessarily diagnostic), we also found clear allometry-independent shape variation. This allowed us to define new, easily measured diagnostic traits in the palate, which differentiate the three clades. Contrary to previous suggestions, we found no support for a latitudinal gradient as causing the differentiation between the clades. However, skull shape co-varied with temperature and precipitation seasonality, suggesting that the clades may be adapted to environmental variables that are likely to be impacted by climate change. Our study demonstrates the use of 3D geometric morphometrics to improve taxonomic diagnosis of cryptic mammalian species, while providing perspectives on the adaptive origins and potential future threats of mammalian diversity.

## I. Introduction

Mammalian biodiversity is globally under threat due to anthropogenic impacts, which include changing patterns of temperature and rainfall, and increased frequency and duration of extreme bushfires (Bowman *et al.*, 2020). In terms of species loss, the Australian mammalian fauna is globally the most affected due to widespread environmental degradation (Woinarski, Burbidge, & Harrison, 2015) from introduced species, agriculture, logging and extreme vegetation fire events (Pardon *et al.*, 2003; Letnic, Tamayo, & Dickman, 2005; Firth *et al.*, 2010; Pastro, 2013; Crowther *et al.*, 2018; Radford *et al.*, 2018; Murphy *et al.*, 2019). Furthermore, Australia is predicted to endure increasingly frequent fire-weather and extreme droughts (Di Virgilio *et al.*, 2019; Dowdy *et al.*, 2019; Ukkola *et al.*, 2020; Kirono *et al.*, 2020). The east coast of Australia is particularly vulnerable in terms of biodiversity loss: the 2019-2020 mega-fires of south-eastern Australia destroyed habitat within the distribution of 832 vertebrate taxa, of which 83 were mammalian species (Ward *et al.*, 2020). Hence, the implementation of widespread conservation efforts for over one hundred threatened Australian mammal species is a high priority (Legge *et al.*, 2018).

The planning of such conservation efforts relies on a good understanding of what species inhabit the most affected environments. However, the advent of modern molecular methods is driving the discovery of previously unrecognized “cryptic” species and is increasingly pointing towards unexpectedly high biodiversity losses, with taxa at risk of extinction shortly after, or even before, discovery (May, 1988; Dubois, 2003). This particularly affects many species that are considered to be “known” but may eventually undergo taxonomic revision with the development of more accurate taxonomic methods (Bickford *et al.*, 2007; Chaplin *et al.*, 2020). Thus, we may be underestimating well-studied species initially classified as not threatened that may instead contain evolutionary lineages sufficiently distinct to deserve re-examination of their conservation status.

The issue of unrecognised biodiversity goes beyond the fundamental question of how many species exist. Characterising the phenotypic diversity in closely related species is also essential for understanding their interaction with the environment – that molecular data alone does not provide – and the reason for the species divergence. However, the phenotypic diversity in young species is generally measured using well-established morphological diagnostics (e.g., Baker, Mutton, & Van Dyck, 2012; Baker *et al.*, 2012; Baker, Mutton, & Hines, 2013) that were designed with a view to species differentiation, rather than the processes that may have led to phenotypic divergence. In order to identify conservation units worthy of protection, we need to understand the ecological processes that may lead to species diversification – in fact, ecological exchangeability is a defining factor of Evolutionary Significant Units (Crandall *et al.*, 2000; Fraser & Bernatchez, 2001; de Guia & Saitoh, 2007). In particular, population divergence and speciation is widely known to occur in association with certain ecological boundaries. For example, in coastal eastern Australia, the Brisbane Valley represents a biogeographic break (Bryant & Krosch, 2016) for divergence of arthropods (Lucky, 2011; Rix & Harvey, 2012), reptiles (Chapple, Chapple, & Thompson, 2011a; Chapple *et al.*, 2011b), amphibians (McGuigan *et al.*, 1998; James & Moritz, 2000) and mammals (Bryant & Fuller, 2014); the Clarence River Corridor (Bryant & Krosch, 2016) is also known for the divergence of reptiles (Colgan, O’Meally, & Sadlier, 2010) and mammals (Rowe *et al.*, 2012; Frankham, Handasyde, & Eldridge, 2012). Thus, assessing the ecological interaction and the distribution of morphologically diversified vertebrate taxa, particularly across biogeographic breaks, is key to understanding the role of biotic and abiotic factors in the divergence of these species.

The marsupial mammal genus *Antechinus* (MacLeay, 1841) is a case in point of unrecognised diversity located on the Australian east coast. Antechinuses are small, scansorial and insectivorous marsupials (Lee & Cockburn, 1985), which play an important role as pollinators (Goldingay, Carthew, & Whelan, 1991; Goldingay, 2000). The genus also displays the unusual trait of semelparity, where all males die after an annual 1-3 week mating period (Kraaijeveld-Smit, Ward, & Temple-Smith, 2003; Holleley *et al.*, 2006; Fisher *et al.*, 2013). Antechinuses are also predicted to be particularly susceptible to changes in rainfall patterns: members of this genus synchronize their only mating event in the life of a male with rainfall-dependant peaks of insect abundance (Fisher *et al.*, 2013).

Antechinus species are a good example of accelerated biodiversity recovery in the wake of recent advances in molecular biology (Baker & Dickman, 2018). Several species in the genus have recently been taxonomically re-described and others have been discovered, expanding their known diversity from 10 to 15 species since 2012 (Baker *et al.*, 2012, 2013, 2014, 2015; Baker & Van Dyck, 2013a,b,c, 2015), with two of these classified as federally Endangered (*Antechinus arktos* and *Antechinus argentus*) (EPBC Act, 1999; Geyle *et al.*, 2018).

The species complex *Antechinus stuartii* has been a focus of taxonomic change since its description by MacLeay in 1841. Five species are currently recognized to have been once part of *A. stuartii* (*sensu lato*): *Antechinus flavipes* (Waterhouse, 1837) (see in Baker & Van Dyck, 2013b); *Antechinus adustus* (Thomas, 1923) (see in Van Dyck & Crowther, 2000); *Antechinus agilis* Dickman *et al.*, 1998, *Antechinus subtropicus* Van Dyck & Crowther, 2000 and *A. stuartii* MacLeay, 1841 (*sensu stricto*; see in Jackson & Groves, 2015). Notably, the difficulties with taxonomic resolution have been driven by a lack of phenotypic differentiation; for instance, *A*. *stuartii* and *A. agilis* were still thought to be morphologically cryptic until near the end of the century (Sumner & Dickman, 1998). Further taxonomic clarification of *A. stuartii* has recently been recommended after genetic studies revealed multiple lineages (Mutton *et al.*, 2019).

Previous traditional morphological work (including linear measurements and discrete characters) found a subtle differentiation between *A. subtropicus* and *A. stuartii*, particularly in cranial size, rostral proportions and palatal morphology (Van Dyck & Crowther, 2000), although the morphological differences were not clearly defined. However, subsequent molecular work (Mutton *et al.*, 2019) suggested that *A. stuartii* contains two lineages and is paraphyletic: *A. subtropicus* and an *A. stuartii* north clade appear genetically more closely related to the exclusion of the *A. stuartii* south clade (Figure 1). These taxa have apparently arisen from a recent speciation event dated from the Pleistocene (~ 2 Mya) (Mutton *et al.*, 2019). Further morphological evidence is therefore required to assess the taxonomy of the northern and southern *A. stuartii* clades and to examine the relationship of both these taxa with *A. subtropicus*.

**Figure 1:**
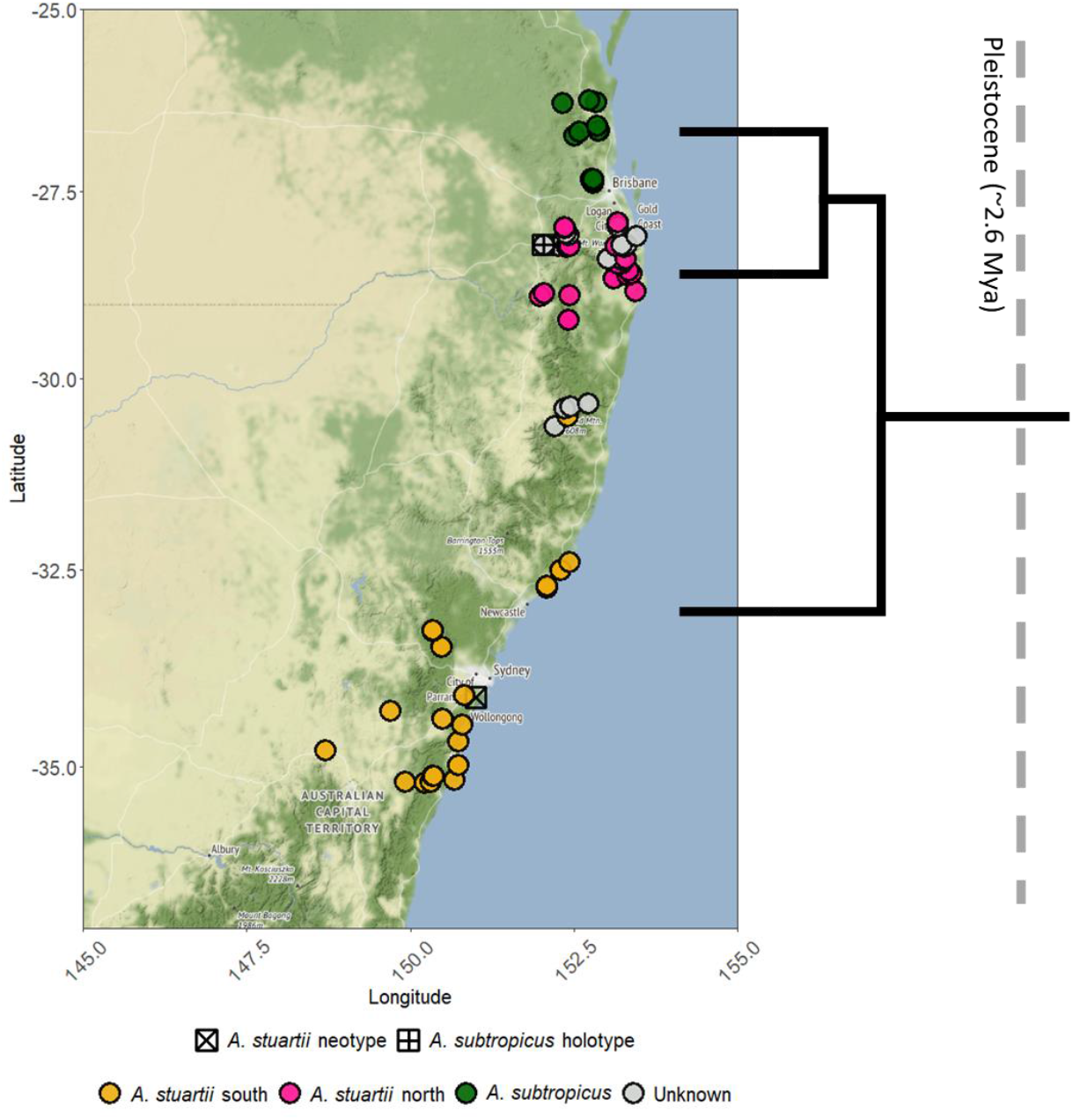
Distribution map of the specimens used for this study. Labelled are A. stuartii south, A. stuartii north, A. subtropicus, specimens of unknown identity within the A. stuartii - A. subtropicus species complex, the holotype of A. subtropicus and the neotype of A. stuartii. All figures in this paper are labelled: A. stuartii south in orange, A. stuartii north in pink and A. subtropicus in green. The phylogeny is adapted from Mutton et al. (2019).

The *Antechinus stuartii* species complex presents a unique opportunity to assess morphological differentiation at the boundaries of a complex of young, closely related species. Because they apparently occur along a latitudinal cline, the *A. stuartii* complex also provides the context for an assessment of ecological and geographic factors that drive species differentiation and are the chief predictors of future species distributions. In this study, we take advantage of 3D geometric morphometrics to investigate the morphological diversification within the *A. stuartii – A. subtropicus* complex. The benefit of geometric morphometrics over conventional morphological measurements in this context is that it allows a global assessment of shape retained through variation and differentiation analyses, with graphical depictions permitting accurate biological interpretations (Adams, Rohlf, & Slice, 2013). We therefore aim to: a) test for corroboration of the genetically known clades with dependable morphological differentiators (taxonomic aspect); b) evaluate the environmental drivers associated within and between clades (ecomorphological aspect); and c) infer an evolutionary hypothesis for the speciation events (evolutionary aspect).

## II. Material and methods

### a) Data collection

Our study included 168 3D models of adult individuals (determined by complete P3 premolar tooth eruption). These included specimens of *Antechinus subtropicus* (n = 68; 41 males, 25 females and 2 of unknown sex), *Antechinus stuartii* north (n = 30; 16 males, 12 females and 2 of unknown sex), *Antechinus stuartii* south (n = 38; 15 males, 22 females and 1 of unknown sex) and *A. subtropicus / A. stuartii* where clades were unassigned (n = 32; 19 males, 8 females and 5 of unknown sex). To determine clade groupings, first, specimens were assigned a species (*A. subtropicus* or *A. stuartii*) following identification by museum curators. Second, we corroborated this information with specimens that were assigned a clade genetically in the literature. Third, we assigned the remaining specimens following the presumed distribution of the clades, as they are largely geographically non-overlapping (Mutton *et al.*, 2019). To assign *A. stuartii* specimens to the northern or southern lineage, we relied on the locality in which they were captured (the lineages are largely geographically non-overlapping, with a narrow zone of sympatry) and genetic analyses (two mtDNA and four autosomal nuclear genes) of population representatives sourced from Mutton et al. (2019). We left a range of specimens unassigned when their source populations were not genetically determined or when they were not genetically determined and were captured at localities near the narrow clades overlap zones. Because of the current lack of clarity around whether the genetic lineages of *A. stuartii* “north” and “south” represent true species, we refer to the three taxonomic units of *A. stuartii* “north”, *A. stuartii* “south”, and *A. subtropicus* as “clades” throughout this manuscript.

We used a GoMeasure 3D HDI109 blue light surface scanner (LMI Technologies Inc. Vancouver, Canada) to create the 3D models. This scanner was portable, allowing us to carry it to museum collections around Australia (Queensland Museum, Australian Museum and Australian National Wildlife Collection). Scanning was undertaken according to protocols developed by Marcy *et al*. (2018) and Viacava *et al*. (2020). We placed each skull specimen in 3 different orientations on a motorised rotary table turning every 45 degrees (8 rotations per round). We meshed together the 24 resulting 3D models with the scanner’s software FLEXSCAN3D 3.3 to produce a final 3D model of the complete skull. This 3D model was then cleaned, decimated and reformatted following Viacava *et al.’s* (2020) protocol. Photographs of each specimen helped identify landmarks – for example, we were able to discriminate the nasal-maxillary suture, visible in photographs, from non-biological 3D artefacts.

The landmarking template is an adaptation of Viacava *et al.’s* (2020) template based on another dasyurid, the northern quoll (*Dasyurus hallucatus*). This template consists of 412 landmarks: 82 fixed landmarks, 63 curves (185 semilandmarks) and 9 surface patches (145 semilandmarks) (see landmark locations and their anatomical definitions in Supplementary Figure 5 and Supplementary Table 1, respectively). We avoided landmarking the zygomatic arches because their thin geometry was not captured well at the scanner’s resolution. Moreover, the preparation of the skeleton can cause errors on small specimens because zygomatic arches can warp after losing support from the muscles during dehydration (Yezerinac, Lougheed, & Handford, 1992; Schmidt *et al.*, 2010). However, the acquisition of a substantial number of specimens and generous coverage of the anatomical zones surrounding the zygomatic arches have been shown to surmount this issue (Marcy *et al.*, 2018).

All fixed landmarks and curves were manually registered by P. V. in Viewbox version 4.0 (dHAL software, Kifissia, Greece). Curve semilandmarks were projected onto the curves, placed equidistantly and were finally slid along their respective curves in Viewbox. Surface semilandmarks followed a thin-plate spline interpolation between the template and each specimen, then were projected to the surface and were finally slid. All sliding procedures were performed following minimization of the bending energy (Bookstein, 1997).

### b) Analyses

We analysed the 3D raw coordinates in R version 3.6.3 (R Core Team, 2019), using the packages “geomorph” (version 3.1.3) (Adams & Otárola-Castillo, 2013), “Morpho” (version 2.7) (Schlager *et al.*, 2019) and “landvR” (version 0.5) (Guillerme & Weisbecker, 2019). The first step was to translate, rotate and scale all specimens to the same size by performing a Generalized Procrustes Analysis (GPA). This method results in the decomposition of centroid size and isometry-free shape variation for further analyses (Rohlf & Slice, 1990). Thus, shape, as defined by Kendall (1989), is the resultant of the form of the object minus its isometric component. To analyse shape in each Kendall’s morphospace, we performed this GPA step for all specimens and also for the corresponding subsets. For example, when only specimens of known sex or group were considered, the superimposed dataset changed by leaving aside those specimens that were unidentified by sex or group. Note that all analyses involving permutations were set to 1000 iterations, and that Bonferroni adjustments of *p*-values were used to correct for tests involving multiple comparisons.

#### Size, allometry and sexual dimorphism

We performed pairwise comparisons between the centroid size least squares means of each clade to assess if the skulls of each clade were differently sized. Note that centroid sizes were within the same order of magnitude, such that log-transformation was not deemed necessary.

We evaluated the influence of size on shape (allometry – the component of shape that depends on size but is not isometric) in the entire dataset with a Procrustes ANOVA. We then used a Homogeneity of Slopes Test to assess whether the allometric slopes differed between sexes and clades. If this was not the case and they shared a common allometric slope, this would enable us to evaluate allometry-free (i.e. free of shape patterns associated to allometry) shape differences between sexes and clades. The latter step requires additional Procrustes ANOVAs including size as a first and sex or clade as a second predictor variable (Table 1).

**Table 1:** ANOVA on predictors of size variation and Procrustes ANOVA on predictors of shape variation.

| RESPONSE VARIABLE | PREDICTOR VARIABLE | QUESTION | D. F. | SS | R<sup>2</sup> | F | Pr(>F) | INTERPRETATION |
| --- | --- | --- | --- | --- | --- | --- | --- | --- |
| Size | Clade | Are clades different in size? | 2 | 5104.8 | 0.338 | 33.925 | 0.001 | Clear effect. |
|  | Sex | Are sexes different in size? | 1 | 5184.7 | 0.352 | 71.46 | < 0.001 | Clear effect. |
| Shape | Clade | Are clades different in shape? | 2 | 0.017 | 0.143 | 11.082 | 0.001 | Clear effect. |
|  | Size | Is there allometry? | 1 | 0.016 | 0.133 | 20.477 | 0.001 | Clear effect. |
|  | Sex | Are sexes different in shape? | 1 | 0.004 | 0.038 | 5.051 | 0.001 | Low effect sizes and low variance explained. |
|  | Size : Sex | As there is sexual dimorphism and allometry, do sexes differ in allometric slopes? | 1 | 0.001 | 0.006 | 0.907 | 0.587 | No clear effect. |
|  | Size + Sex | Adjusting for size, are sexes different in shape? | 1 | 0.001 | 0.013 | 1.946 | 0.016 | Low effect sizes and low variance explained. |
|  | Size : Clade | Do clades differ in allometric slopes? | 2 | 0.001 | 0.012 | 1.014 | 0.433 | No clear effect. |
|  | Size + Clade | Adjusting for size, are clades different in shape? | 2 | 0.01 | 0.085 | 7.175 | 0.001 | Clear effect. |

#### Inter- and intra-group shape variation

To explore the patterns of the main shape variation and to assess whether it corresponded to clade boundaries, we conducted a Principal Component Analysis (PCA) on the landmark coordinates. We labelled the groups in the plot of the First and Second Principal Component (PC) scores. We then conducted a Procrustes ANOVA to test whether the groups were morphologically different. We then performed permutation-based pairwise comparisons between mean shapes of these groups.

We also performed pairwise comparisons of disparity to evaluate if the clades and sexes differed in morphospace occupancy. Several measurements of disparity are possible (Guillerme *et al.*, 2020), but here we consider the widely-used Procrustes variance. This asks if the residuals of a common linear model fit differ in magnitude between groups, which is possible even if there are no significant differences in shape between groups.

Additionally, we aimed to determine the linear distances between landmarks that our analyses suggested to be the most variable between clades. The linear measurements that best distinguish the clades were determined by creating heat maps of landmark displacement between the mean shapes of each clade (which is also what is used to determine statistical differences between clades in Procrustes ANOVAs). We colourised landmarks according to how great the displacement is between shapes for each landmark (see the “landvR” package for details) (Guillerme & Weisbecker, 2019). The darkest, most displaced among the landmarks were chosen as candidates for linear measurements to differentiate clades. The goal was to provide a measure of shape differentiation between clades that was easily reproducible in taxonomic museum work. To test for differences between clades, we ran a linear model of these linear distances against centroid size to correct for size, and performed post hoc Tukey multiple comparison tests between means of the clades.

#### Ecomorphological characterisation

We conducted a variation partitioning analysis of cranial shape for variables that potentially contribute to cranial shape variation. The factors included in the final model were size (as centroid size), the geographic distances among specimens (based on location of capture) and four environmental variables (temperature, temperature seasonality, precipitation and precipitation seasonality). Elevation was initially included as a geographic factor but did not have a clear effect on shape variation and was therefore not retained in the final model. We avoided spatial autocorrelation by transforming the latitude and longitude coordinates of each individual into a principal coordinates neighbourhood matrix (Borcard & Legendre, 2002) and retaining only the axes with positive eigenvalues. We tested for significance in shape variation of the selected axes and included those that were significant in the final variation partitioning model of shape variation. We obtained the environmental variables from the Atlas of Living Australia (www.ala.org.au) and WORLDCLIM (v 2.0) (www.worldclim.org/bioclim) (O’Donnell & Ignizio, 2012). Temperature seasonality (BIO4) is calculated as the standard deviation of the monthly mean temperatures to the mean monthly temperature. Precipitation seasonality (BIO15) is calculated as the ratio of the standard deviation of the monthly total precipitation to the mean monthly total precipitation. Both seasonality variables are known as coefficients of variation and are expressed as a percentage. To partition the shape variation with respect to these environmental variables, we used the *varpart* function in the “vegan” package for R version 3.6.3 (Oksanen *et al.*, 2018). We complemented this analysis with a redundancy analysis ordination on partial and full models (1000 permutations). Finally, to discern how each environmental variable influences shape, we performed a separate variation partitioning analysis of cranial shape with only the four environmental variables.

## III. Results

### Allometry and sexual dimorphism

The statistics for all the following results are shown in Table 1. In the entire dataset, Procrustes ANOVA revealed that size accounted for 13.3 % of the shape variation (R^2^ = 0.133; p = 0.001). Males were significantly larger than females (also see boxplots in Supplementary Figure 1). Without allometric correction, we found small shape differences between sexes. However, both sexes followed the same allometric slope, allowing us to test if the sexual shape differences were only due to the differential size between sexes. After accounting for size, no clear differences were found between sexes. Thus, size differences between males and females almost entirely account for the shape differences between sexes.

Centroid size differences were clear only between the larger *A. subtropicus* and the smaller *A. stuartii* (see also boxplots in Figure 2). Before allometric correction, 14.3% of shape variation of the sample was associated with shape differentiation between clades. As with sexes, the clades followed a common allometric slope allowing us to test if the shape differences between clades were purely due to allometry. Unlike the sex comparisons, this was not the case: allometry-free shape differences between clades were significant and accounted for 8.5% of the shape variation.

**Figure 2:**
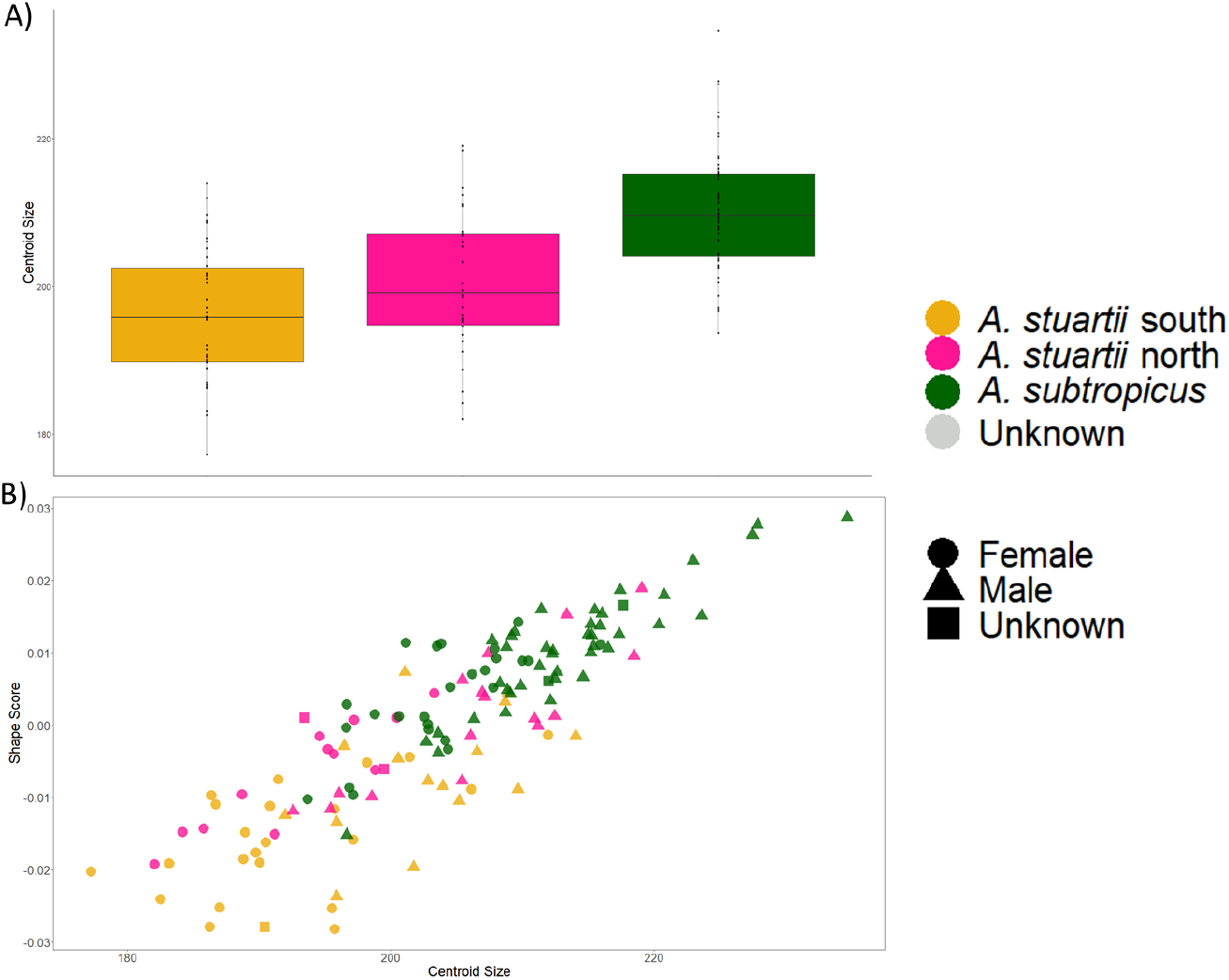
A) Box plot and dot plot of centroid size labelling each clade as per Figure 1. Centroid size differences were clear only between the *larger A. subtropicus* and the smaller *A. stuartii* (both mean comparisons between *A. subtropicus* and the two clades of *A. stuartii* were significant; p = 0.003), but not between *A. stuartii* south and *A. stuartii* north (p = 0.282). B) Allometry plot consisting of centroid sizes versus shape scores obtained from the regression of shape on size (Drake & Klingenberg, 2008).

### Inter- and intra-group shape variation

The first two principal components represented 28.23 % of the total shape variation (PC1 = 19.68%; PC2 = 8.55%) (Supplementary Figure 2). We found clear shape differences between clades (R^2^ = 0.143; F_1, 127_ = 11.082; p = 0.001). Pairwise comparisons showed that the three clades clearly differed in shape from each other (all three pairwise comparisons between means: p = 0.001) (also see Figure 3). *Antechinus subtropicus* had larger major palatine foramina and larger incisive foramina, thus with a smaller interpalatal distance compared to both *A. stuartii* lineages. *Antechinus stuartii* north had larger major palatine foramina than *A. stuartii* south and smaller incisive foramina than *A. subtropicus* (similar to *A. stuartii* south). *Antechinus stuartii* south had smaller major palatine foramina than *A. stuartii* north and *A*. *subtropicus*. We also observed a slight difference of molar row length, larger in *A. stuartii* south when compared to *A. stuartii* north and *A. subtropicus*. We found no clear morphological disparity (Procrustes variance) differences between lineages (*A. stuartii* south vs *A. stuartii north*, p = 0.712; *A. stuartii* north vs *A. subtropicus*, p = 0.93; *A. stuartii* south vs *A. subtropicus*, p = 0.734).

**Figure 3:**
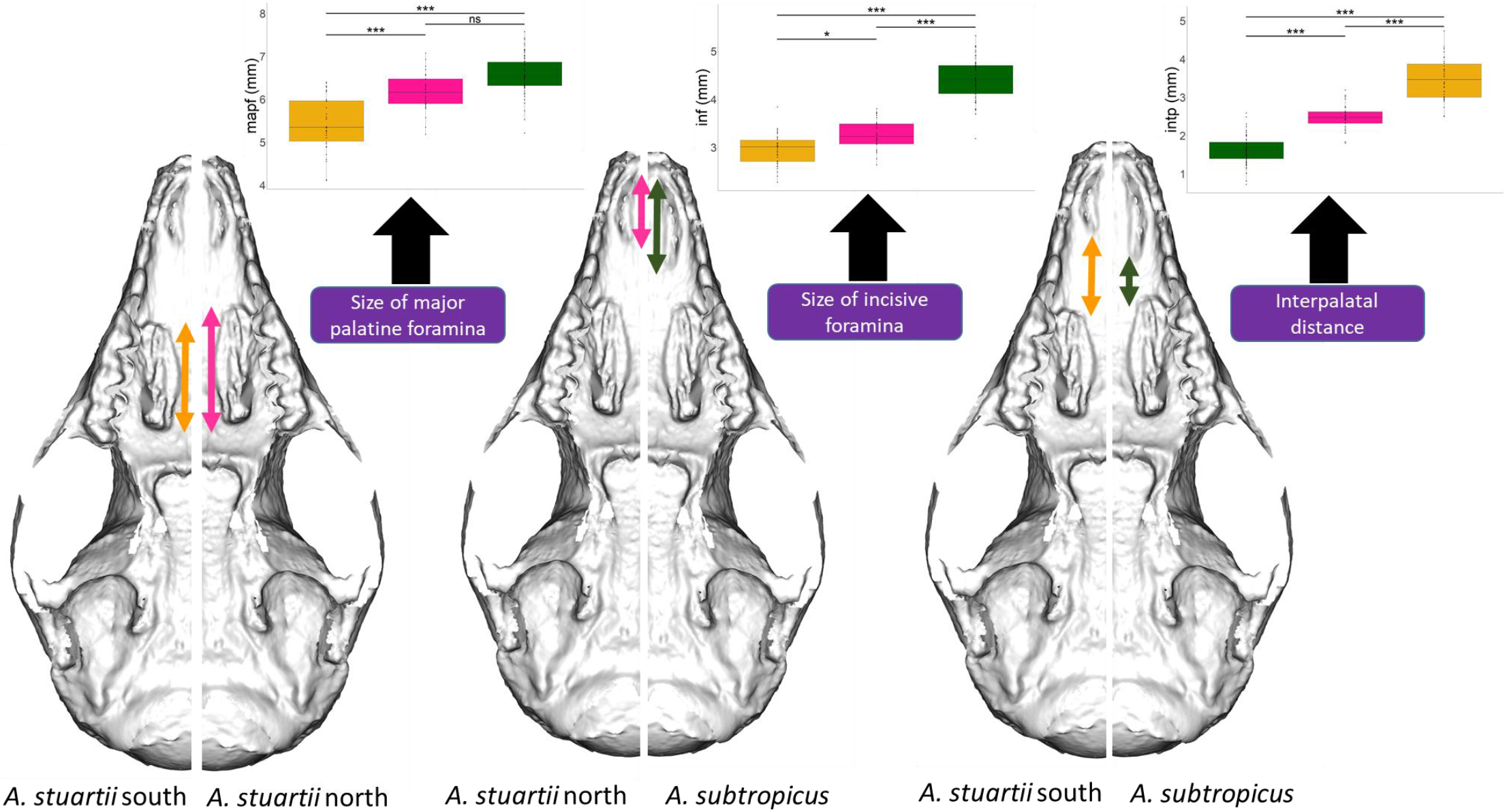
Pairwise comparisons between mean shapes of each clade (*A. stuartii* south vs *A. stuartii* north, p = 0.003; *A. stuartii* north vs *A. subtropicus*, p = 0.003; *A. stuartii* south vs *A. subtropicus*, p = 0.003; all p-values were adjusted with following the Bonferroni method). The 3D images are the specimen closest to the overall mean warped correspondingly to the mean shapes of each clade. Tukey post-hoc analyses of linear measurements after size correction were performed; significance levels (*p<0.05, **p<0.01, ***p<0.001) are shown in the boxplots. For each comparison, we label the best differentiator diagnostic; i.e., the size of the major palatine foramina (mapf) for differentiating *A. stuartii* south and *A. stuartii* north, the size of the incisive foramina (inf) for differentiating *A. stuartii* north and *A. subtropicus*, and the interpalatal distance (intp) for differentiating between the three clades. Clades are consistently labelled as per Figure 1.

Heat maps of landmark displacements between clade mean shapes revealed a striking dominance of just a few landmarks differentiating them. These were the most anterior point of the major palatine foramen and the most posterior point of the incisive foramen (Supplementary Figure 3). We therefore deemed the most distinguishable character between the three lineages to be the distance between the major palatine and incisive foramen (the interpalatal distance – Fig. 3). We measured this distance, averaged both sides for each specimen and determined the distance ranges between these two landmarks for each lineage (Figure 3). All three lineages were clearly different in this character between each other (p < 0.001). Post hoc Tukey’s multiple comparisons between clade means of three linear distances involving the most variable landmarks revealed linear measurements with potential for distinguishing clades (see boxplots of Figure 3). The size of the major palatine foramina differentiated *A. stuartii* south vs *A. stuartii* north (p < 0.001) and *A. stuartii* south vs *A*. *subtropicus* (p < 0.001) but did not differentiate *A. stuartii* north vs *A. subtropicus* (p = 0.678). The size of the incisive foramina differentiated *A. stuartii* south vs *A. subtropicus* (p < 0.001), *A. stuartii* north vs *A. subtropicus* (p < 0.001) and *A. stuartii* south vs *A. stuartii* north (p = 0.0228). The interpalatal distance differentiated *A. stuartii* south vs *A. subtropicus* (p < 0.001), *A. stuartii* north vs *A. subtropicus* (p < 0.001) and *A. stuartii* south vs *A. stuartii* north (p < 0.001).

### Ecomorphological characterization

The geographic effect on shape was significant in the *A. stuartii - A. subtropicus* species complex (refer to Table 2 for significance levels and effect sizes). Latitude and longitude were significantly contributing factors to both shape and size variation (see Table 2). However, the geographic variation is confounded with the geographic distribution of clades along the east coast: within-clade geographic analyses did not show clear effects latitudinally nor longitudinally on size and shape (Table 2). Only members of *A. stuartii* south showed significant geographic variation (latitude and longitude) in shape, but they were weakly related (Table 2).

**Table 2:** Analyses of Variance on geographic sources of size and shape variation of the entire A. stuartii - A. subtropicus species complex and within each clade.

| RESPONSE VARIABLE | PREDICTOR VARIABLE | QUESTION | SS | R<sup>2</sup> | F | Pr (>F) | INTERPRETATION |
| --- | --- | --- | --- | --- | --- | --- | --- |
| Size | Latitude | Is latitude covarying with size in this dataset? | 3735.5 | 0.195 | 40.29 | 0.001 | Clear effect. |
|  | Latitude within each clade | Is latitude covarying with size within each clade? | South: 104.32<br>North: 303.04<br>Sub: 4.7 | South: 0.035<br>North: 0.111<br>Sub: 0.001 | South: 1.311<br>North: 1.08<br>Sub: 0.073 | South: 0.261<br>North: 0.061<br>Sub: 0.813 | No clear effect. |
|  | Longitude | Is longitude covarying with size in this dataset? | 3428.9 | 0.179 | 36.261 | 0.001 | Clear effect. |
|  | Longitude within each clade | Is longitude covarying with size within each clade? | South: 21.16<br>North: 485.53<br>Sub: 31 | South: 0.007<br>North: 0.178<br>Sub: 0.007 | South: 0.258<br>North: 6.07<br>Sub: 0.478 | South: 0.629<br>North: 0.027<br>Sub: 0.461 | No clear effect. North might have a biased sample. |
| Shape | Latitude | Is latitude covarying with shape in this dataset? | 0.014 | 0.093 | 17.054 | 0.001 | Clear effect |
|  | Latitude within each clade | Is latitude covarying with shape within each clade? | South: 0.002<br>North: 0.001<br>Sub: 0.001 | South: 0.07<br>North: 0.052<br>Sub: 0.025 | South: 2.705<br>North: 1.543<br>Sub: 1.667 | South: 0.001<br>North: 0.058<br>Sub: 0.039 | Only south stuartii is varying latitudinally in shape with low effect. |
|  | Longitude | Is longitude covarying with shape in this dataset? | 0.012 | 0.081 | 14.645 | 0.001 | Clear effect |
|  | Longitude within each clade | Is longitude covarying with shape within each clade? | South: 0.002<br>North: 0.001<br>Sub: 0.001 | South: 0.066<br>North: 0.051<br>Sub: 0.022 | South: 2.554<br>North: 1.494<br>Sub: 1.461 | South: 0.003<br>North: 0.09<br>Sub: 0.117 | Only south stuartii is varying longitudinally in shape with low effect. |

We partitioned the contribution of geography, size and climate (precipitation + precipitation seasonality + temperature + temperature seasonality) to the varpart model (Figure 4). The full model showed a significant effect on shape variation (F_15,152_ = 3.981, adjusted R^2^ = 0.211, p = 0.001). Pure geographic distances contributed 3% on shape variation (F_10,152_ = 1.593, adjusted R^2^ = 0.029, p = 0.001). Size alone explained 8% of the variance in the varpart model (F_1,152_ = 17.025, adjusted R^2^ = 0.083, p = 0.001). Climatic variables alone only contributed to less than 1% of the variance (F_4,152_ = 1.375, adjusted R^2^ = 0.008); however, when considered jointly with geography, climate and geography explained 9% of the shape variation in the model (F_4,163_ = 5.211, adjusted R^2^ = 0.092, p = 0.001). Of the four environmental variables considered in the model, precipitation seasonality and temperature jointly contributed the most to shape variation (Figure 4 and Table 3).

**Figure 4:**
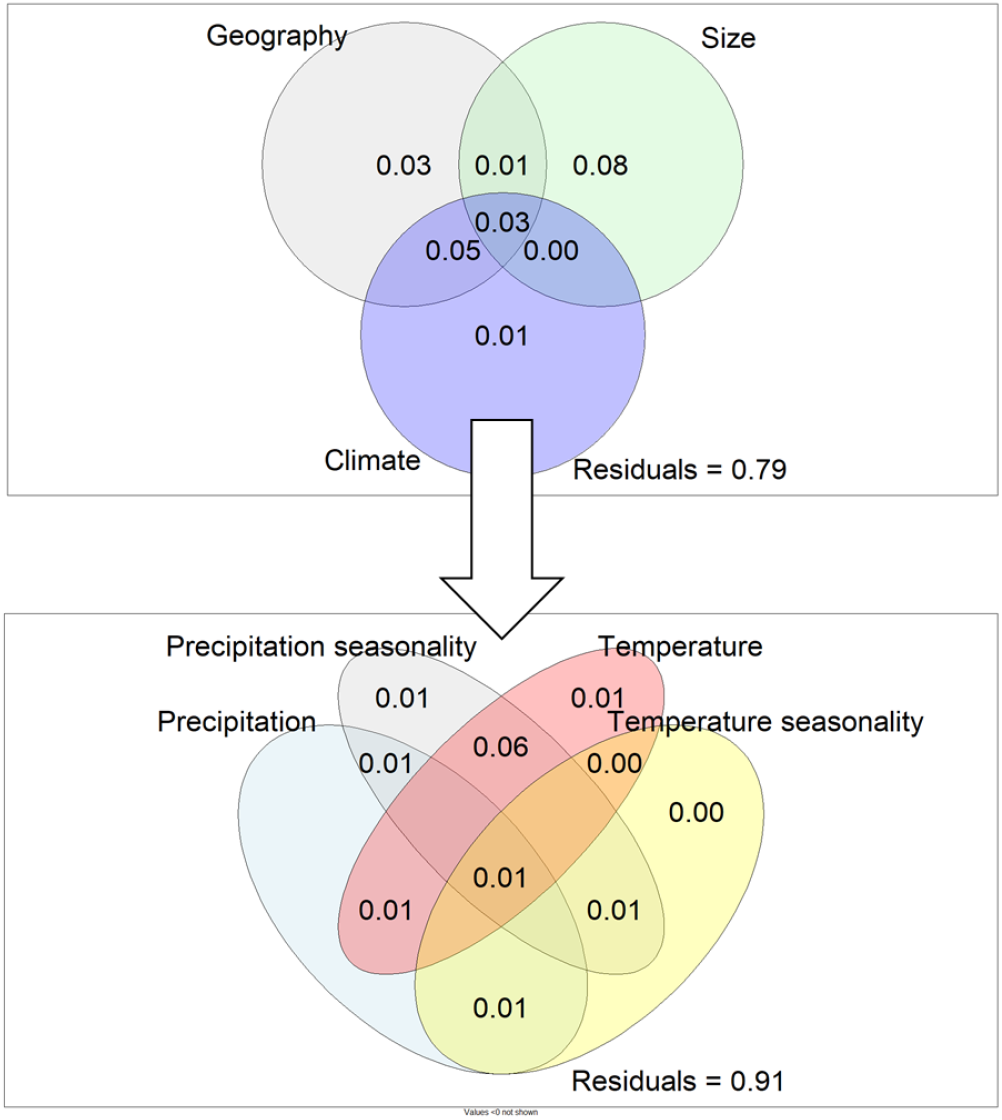
Venn diagrams illustrating variation partitioning analyses. Each individual fraction for each factor contributing to the model is shown in every set. Circle sizes and white space out of the circles representing the unexplained variation are schematic and not to scale.

**Table 3:** ANOVA on climatic predictors of size variation and Procrustes ANOVA on climatic predictors of shape variation.

|  | d. f. | Size |  |  |  | Shape |  |  |  |
| --- | --- | --- | --- | --- | --- | --- | --- | --- | --- |
|  |  | SS | R<sup>2</sup> | F | Pr(>F) | SS | R<sup>2</sup> | F | Pr(>F) |
| Precipitation | 1 | 731 | 0.032 | 6.596 | 0.011 | 0.002 | 0.013 | 2.135 | 0.012 |
| Precipitation seasonality | 1 | 3193.6 | 0.162 | 33.27 | <0.001 | 0.012 | 0.08 | 14.521 | 0.001 |
| Temperature | 1 | 4550.7 | 0.233 | 51.83 | <0.001 | 0.011 | 0.075 | 13.5 | 0.001 |
| Temperature seasonality | 1 | 1906.7 | 0.094 | 18.38 | <0.001 | 0.002 | 0.016 | 2.69 | 0.004 |
| Elevation | 1 | 390.2 | 0.015 | 3.457 | 0.065 | 0.001 | 0.006 | 1.048 | 0.382 |

## IV. Discussion

*Antechinus* species have undergone multiple taxonomic revisions, with recent genetic data suggesting substantially higher biodiversity within the genus than previously expected. Here, we corroborated the genetic differences between *A. stuartii* north and *A. stuartii* south observing clear morphological differences in the major palatine foramina. Thus, we have provided a cranial morphological differentiator in support of Mutton et al.’s (2019) suggestion that *A. stuartii* south and *A. stuartii* north should be reclassified as separate species, within the scope of the phylogenetic species concept (Nixon & Wheeler, 1990). We also corroborated shape differences between *A. subtropicus* and *A. stuartii* and provided an easy-to-follow morphological differentiation protocol for skulls of the three lineages. Additionally, among the environmental variables considered, precipitation seasonality and temperature were the most important factors for shaping the skull in the *A. stuartii - A. subtropicus* complex. This renders the species group of particular conservation concern because it is located on the east coast of Australia, a zone increasingly impacted by climate change (Di Virgilio *et al.*, 2019; Dowdy *et al.*, 2019; Kirono *et al.*, 2020), which is expected to cause variation in precipitation seasonality, temperature and fire weather. Importantly, the distributional ranges of *A. stuartii* south and *A. stuartii* north fall directly within the burn zone of 2019-2020 mega-fires, and the results of the present study add support to a case for the separate management of these taxa.

Differences in palatal vacuity size and molar row length, which we found as important differentiators in our investigation, are widely recognized as diagnostically useful for species identification in various marsupials. For example, the size of the palatal vacuities differentiate species of potoroids, e.g., *Bettongia* (McDowell *et al.*, 2015) and peramelids, e.g., *Microperoryctes* (Groves & Flannery, 1990). Dasyurid species have also been diagnosed in consideration of palatal vacuity size, e.g., *Sminthopsis* (Archer, 1981; Kemper *et al.*, 2011); and more specifically, several species of *Antechinus* (Van Dyck, 1982; Dickman *et al.*, 1998; Baker *et al.*, 2012, 2014). We also add a new differentiating skull trait of molar row length for the clades studied here, which is possibly linked to allometry. This taxonomic differentiator has been found for other antechinus species varying in absolute size (Baker *et al.*, 2012, 2013; Baker & Van Dyck, 2013b). Molar row length has also been found to separate diverse mammals such as rodents (Anderson & Yates, 2000; Christoff *et al.*, 2000; Gonçalves, Almeida, & Bonvicino, 2005; Boroni *et al.*, 2017), shrews (Balčiauskienė, Juškaitis, & Mažeikytė, 2002), bats (Bogdanowicz, 1990), wombats (Black, 2007) and didelphids (Voss, Lunde, & Jansa, 2005).

Our geographic analyses suggest that the latitudinal shape variation observed across the entire dataset is likely driven by morphological differences between the taxa; however, this latitudinal shape variation is not strongly associated with shape within each taxon (see Table 2). This suggests that clinal variation does co-vary with the shape differentiation between the three clades and the clinal variation we observe in the whole dataset is a result of, but not notably influenced by, the clinal distribution of the three taxa. This appearance of a clinal effect would be even stronger under the old taxonomic combination of *A. stuartii* south and north as one species (rather than *A. stuartii* north being sister taxa of *A. subtropicus*), which may have prompted previous suggestions that the variation between species of *Antechinus* may be driven clinally (Van Dyck & Crowther, 2000) (i.e., climatically/geographically). However, the nature of the variation within the genus and its relation to latitude should be investigated in a broader sample of antechinus species, particularly since there have been suggestions that *A*. *subtropicus* is more similar morphologically to the less genetically closely related *A. agilis* than it is to *A. stuartii* (Dickman *et al.*, 1998; Van Dyck & Crowther, 2000; Crowther, 2002; Crowther, Sumner, & Dickman, 2003). Regardless, finding little or no clinal shape variation *within* each clade (also suggested for the whole of *A. stuartii* by (Crowther *et al.*, 2003)) suggests that speciation-related differences in morphology are independent of within-species variation in these antechinus taxa (a pattern also found in wombats (Weisbecker *et al.*, 2019)).

Our results add some insights on the putative processes behind the speciation event in the young clades of the *A. stuartii – A. subtropicus* complex. In 1981, Archer argued that the species of *Sminthopsis* located in inland arid areas had larger palatal vacuities and linked the size of the vacuities to aridity. We also found differences in vacuity sizes with smaller major palatine foramina in the southern clade of *A. stuartii* relative to the northern clade and *A*. *subtropicus*. Although we did not find a strong influence of precipitation or aridity with shape variation in this species complex, shape variation was strongly associated with temperature and precipitation seasonality variation. In particular, rainfall seasonality is associated with food abundance predictability (Kishimoto-Yamada & Itioka, 2015), which appears tied to variation in breeding times observed in antechinus species and is hypothesized to have driven the evolution of semelparity in these dasyurids (Fisher *et al.*, 2013). This environmental factor may have also influenced reproductive isolation involving morphological differentiation. Intriguingly, the shape changes associated with precipitation seasonality involve differences in the size of the palatal vacuities with *A. subtropicus* displaying larger incisive and major palatine foramina than *A. stuartii*. These vacuities convey access to the vomeronasal organ that plays a major role in antechinus reproduction (Aland, Gosden, & Bradley, 2016).

The difference in molar row length between *A. stuartii* south relative to *A. stuartii* north is noteworthy because it is reminiscent of a well-known effect where carnassial tooth length differentiates carnivoran species living in sympatry. Garcia-Navas et al. (2020) suggested that, due to the mostly insectivorous nature and lack of carnassial teeth in dasyurids, such an effect is not expected; however, the molar row of dasyurids has in the past been argued to act as a single contiguous shearing blade (Werdelin, 1986, 1987; Smits & Evans, 2012). It is therefore possible that the difference in molar row length we observe is related to an effect of dietary niche partitioning, similar to carnivorans. However, because the molar row length differences are tied to allometry, it is also possible that this pattern is influenced by a more general effect of allometric scaling, such as the distinction of molar row length observed between differently sized *A. argentus, A. flavipes* and *A. mysticus* (Baker *et al.*, 2012, 2013; Baker & Van Dyck, 2013b).

Not all of our findings based on geometric morphometrics are consistent with previous morphological observations based on the analysis of skull proportions. For example, the differentiation of *A. stuartii* south with *A. subtropicus* according to molar row length was not observed by Van Dyck & Crowther (2000). Intriguingly also, we did not observe a previously suggested morphological distinction – a longer and narrower rostrum – between *A. subtropicus* and *A. stuartii* (Van Dyck & Crowther, 2000). It is possible that the latter discrepancy is mainly due to pure size differences (i.e., non-shape differences) between these two clades, which may be reflected differently in the different analytical approaches. Specifically, the moderate size-related (allometric) shape changes we observe with geometric morphometrics might not be regarded as allometric in linear measurement studies. Traditional morphometric studies focus on maximal lengths and widths, and use ratios as a descriptor for shape. By comparison, in geometric morphometrics, the descriptor of shape includes the relative positions and distances of all landmarks, and size is removed from the equation after Procrustes superimposition. This can cause two types of discrepancy in the analysis of relative size: first, when allometry is analysed in traditional methods, it generally relies on non-scaled or log-scaled measures, whereas when allometry is analysed in geometric morphometrics, the factors taken into account are the calculated centroid size and a scaled abstract shape (Mitteroecker & Gunz, 2009). Thus, it is possible that rostral length measurements may be heavily influenced by differences in size between species, rendering the diagnostic less suitable for differentiating similarly sized individuals of these clades. A potential second issue is that geometric morphometric landmarking protocols of the mammalian cranium generally rely on homologous points such as suture intersections (type I landmarks), rather than frequently employed taxonomic measures of maximum and minimum widths or lengths. Such “extreme points” of shape may therefore be less emphasised by the landmarking protocol. In such cases, the chief consideration should be whether the location of extreme-point measurements relative to the skull may matter.

In this study, we have demonstrated the versatility of geometric morphometric research, providing taxonomic discernment in otherwise morphologically cryptic species and inferring biological processes by identifying associations between morphological differentiation, and geographic and environmental factors. On the taxonomic aspect, the resolution of these small cryptic taxa by 3D shape analyses highlights the importance of the method in systematic studies. In the future, these mammal taxa may see their geographic ranges reduced by an elevation to species rank and, unfortunately, their populations diminish due to a fatal coincidence with fire line zones along Eastern Australia.

## V. ACKNOWLEDGEMENTS

We thank all museum collections who granted us access to scan their specimens: Heather Janetzki (Queensland Museum), Sandy Ingleby (Australian Museum), and Christopher Wilson and Leo Joseph (Australian National Wildlife Collection from the Commonwealth Scientific and Industrial Research Organisation). P. V. was supported by a University of Queensland Research Training Tuition Scholarship and a University of Queensland Research Higher Degree Living Stipend Scholarship. V. W. was supported by the Future Fellowship FT180100634; and V. W. and M. J. P. were funded by the Australian Research Council Discovery Project DP170103227. We also thank Bastien Mennecart and one anonymous reviewer for their generous comments and helping improve the manuscript.

## VI. AUTHOR CONTRIBUTIONS

P. V. and V. W. conceived the ideas; P. V. collected the data; P. V. and V. W. analysed the data with input from A. M. B., S. P. B. and M. J. P.; and P. V. and V. W. led the writing with input from A. M. B., S. P. B. and M. J. P.

## VII. DATA AVAILABILITY STATEMENT

Data and R code are publicly available in GitHub. 3D models can also be publicly accessed through Morphosource.

