## Supplementary Figures for "Using 3D geometric morphometrics to aid taxonomic and ecological understanding of a recent speciation event within a small Australian marsupial (genus *Antechinus*)"

**Supplementary Figure 1.** Box plot and dot plot of centroid size labelling males and females.

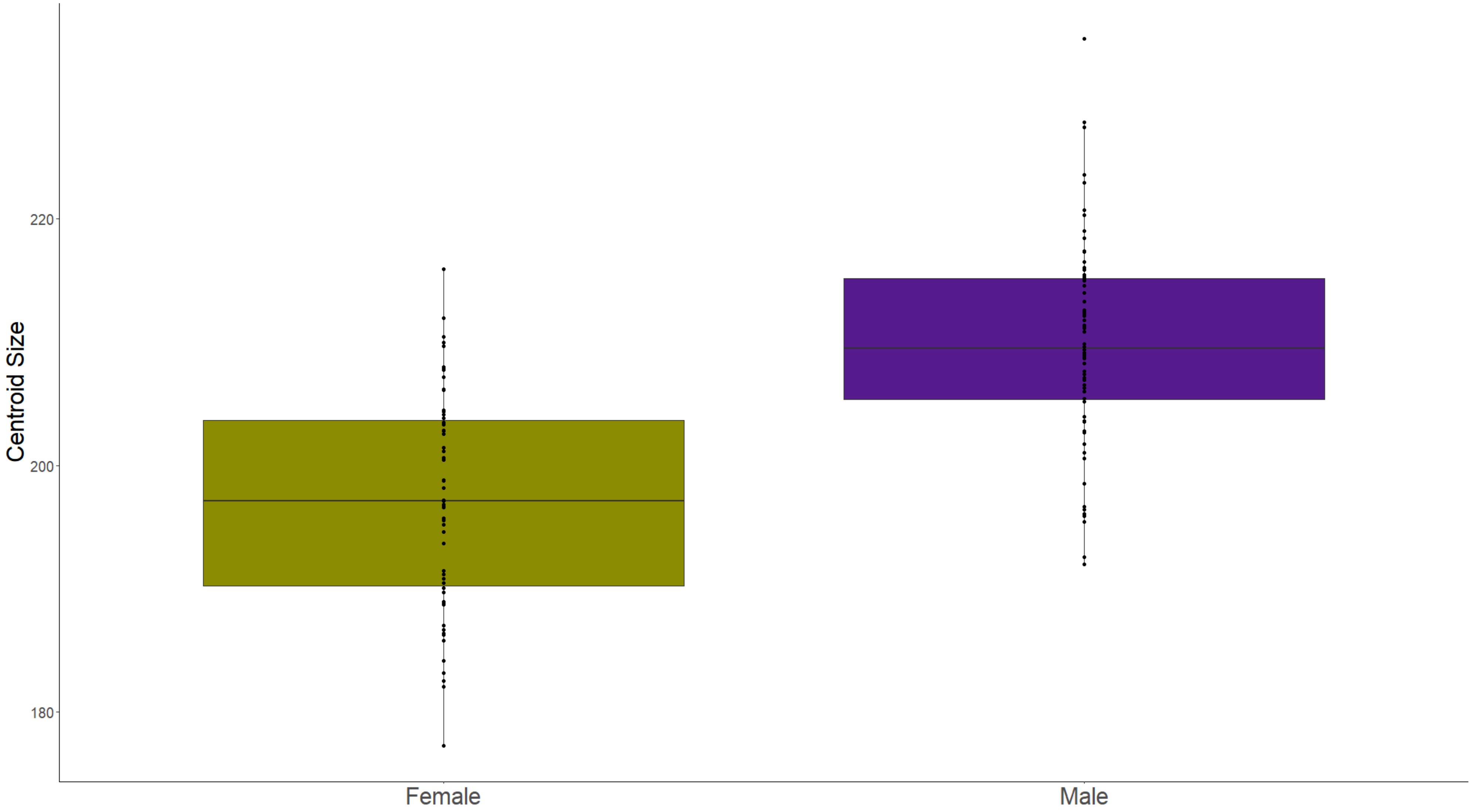
Centroid Size
220
200
180
Female
Male

**Supplementary Figure 2.** Principal Component Analysis on all specimens. Heatmap plots represent the landmark variation from minimum to maximum values of PC1 and PC2 in lateral and ventral views.

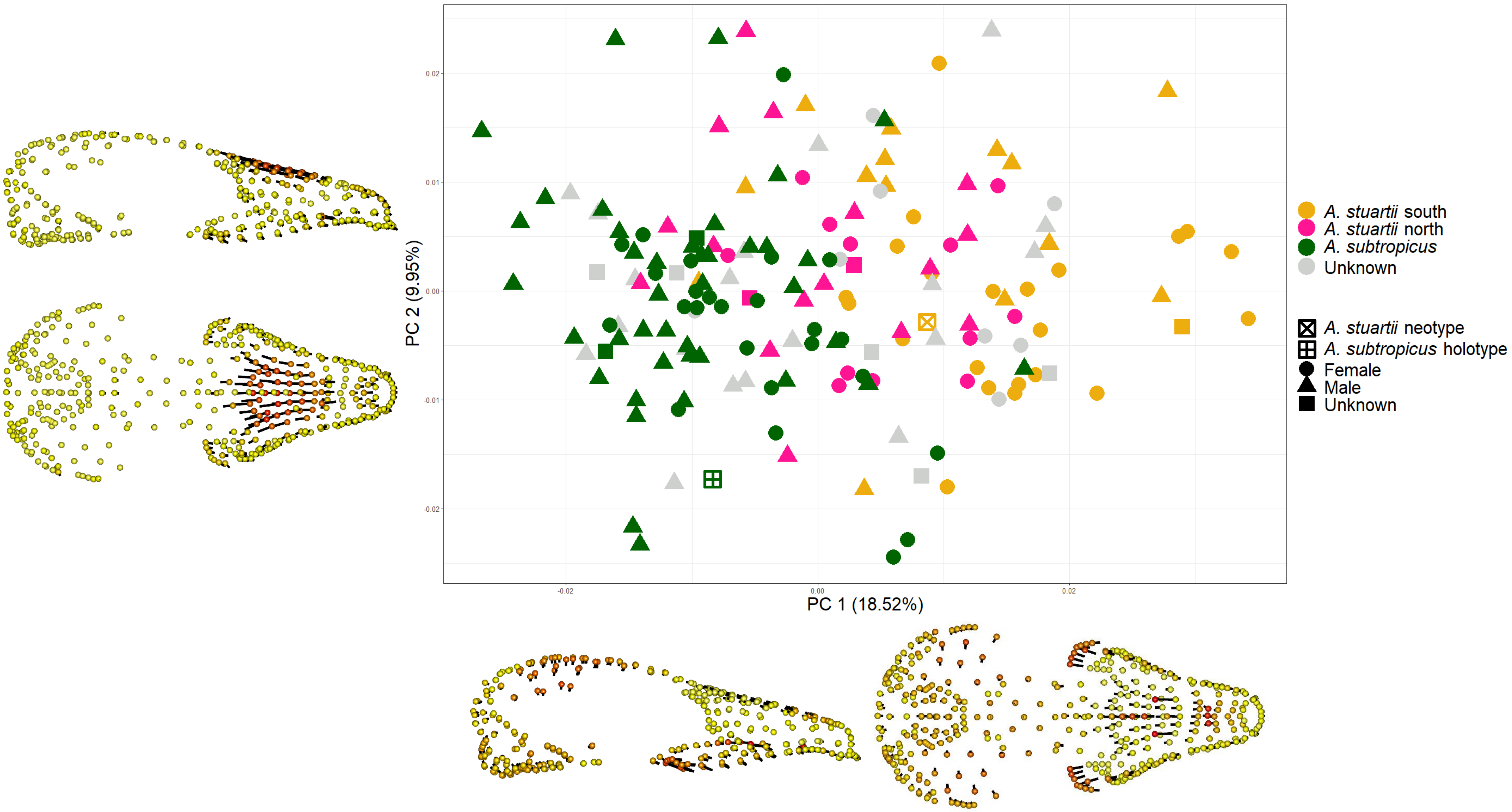
PC 2 (9.95%)
PC 1 (18.52%)
0.02
0.01
0.00
-0.01
-0.02
-0.02
0.00
0.02
A. stuartii south
A. stuartii north
A. subtropicus
Unknown
A. stuartii neotype
A. subtropicus holotype
Female
Male
Unknown

**Supplementary Figure 3.** Heat map plots representing the landmark variation associated to the differences between the mean shapes of *A. stuartii* south, *A. stuartii* north and *A. subtropicus*. Note that the variation is exclusively located in the landmarks of the incisive and major palatine foramina. A slight variation associated to allometry is also observable in the landmarks corresponding to the molar row length.

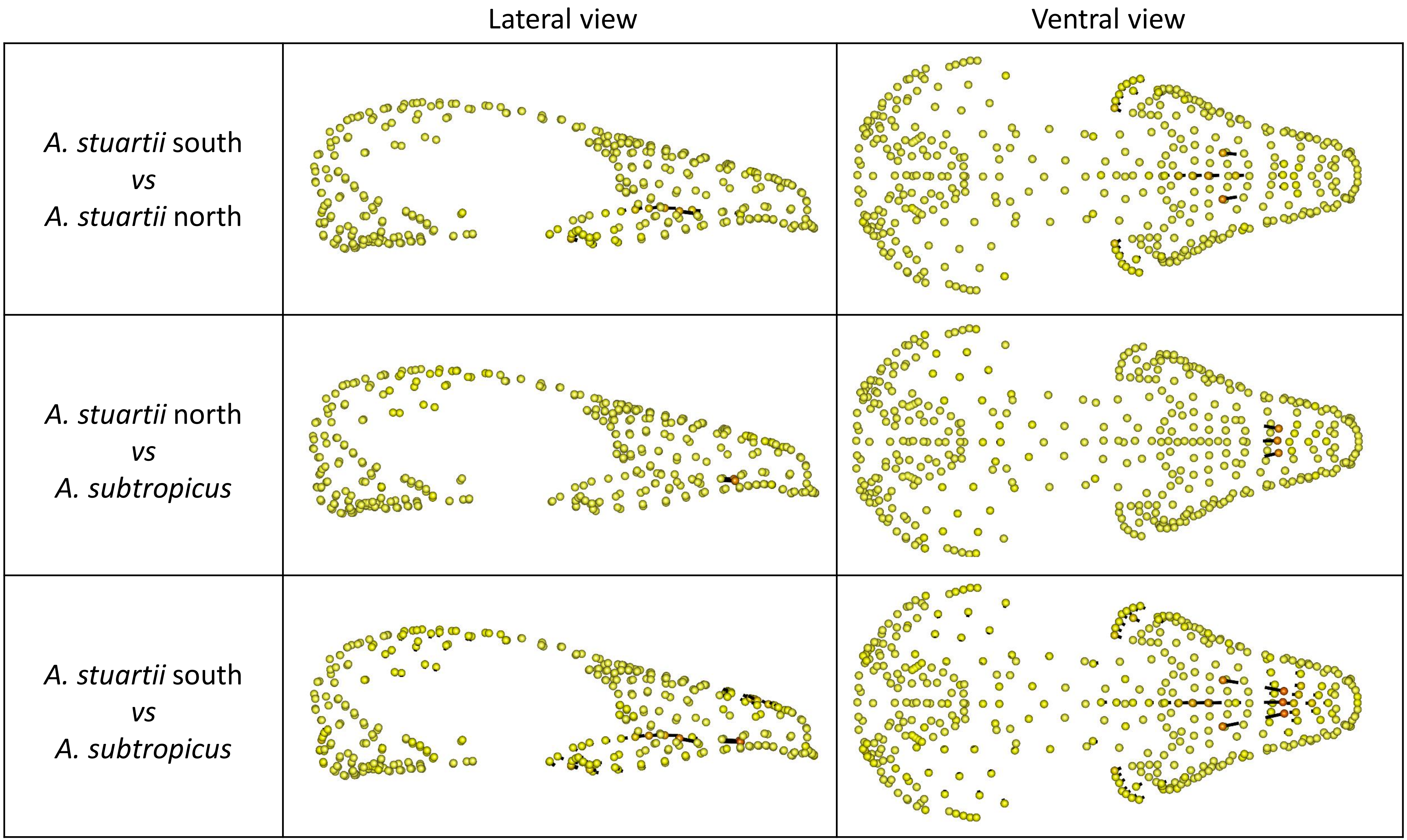

Lateral view
Ventral view
A. stuartii south vs A. stuartii north
A. stuartii north vs A. subtropicus
A. stuartii south vs A. subtropicus

**Supplementary Figure 4.** Heat map plots representing the landmark variation associated to allometry, from minimum to maximum centroid size.

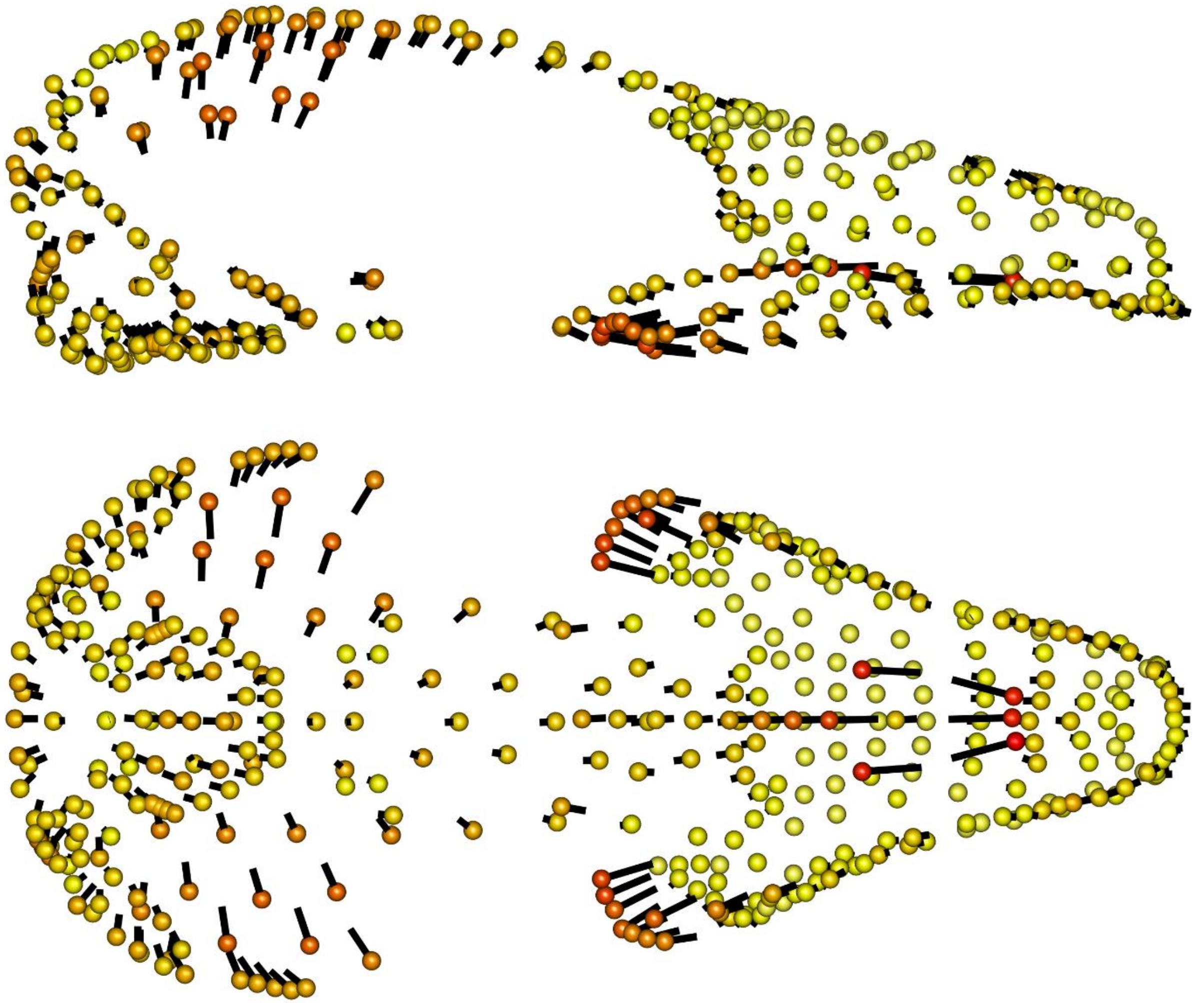

**Supplementary Figure 5.** Template adapted from Viacava et al. (2020) containing 412 landmarks: 82 fixed landmarks (numbered in this Figure), 63 curves (185 semilandmarks) and 9 surface patches (145 semilandmarks). All landmark definitions are described in Supplementary Table I.

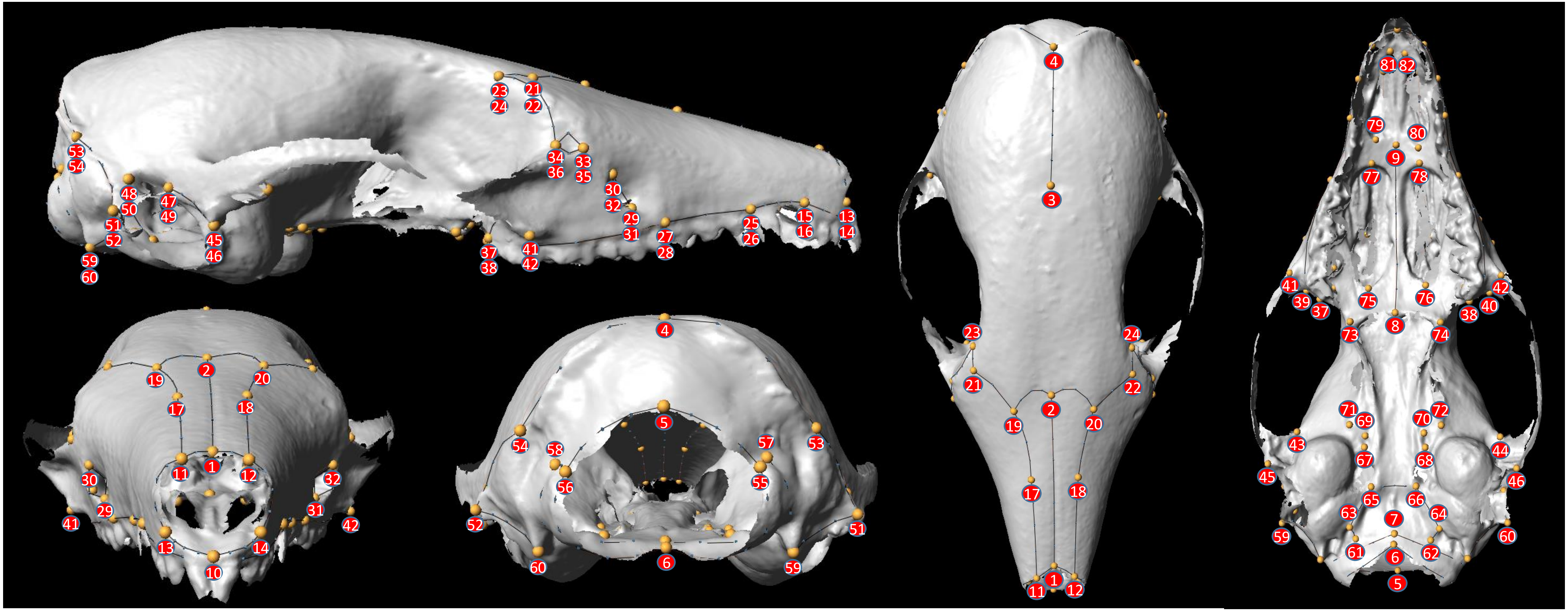
23
21
24
22
53
54
34
33
36
35
48
47
30
50
49
32
51
29
52
45
46
31
59
60
37
38
41
42
27
28
25
26
15
16
13
14
4
3
23
24
21
22
2
19
20
17
18
11
1
12
81
82
79
80
9
77
78
41
42
39
37
75
76
38
40
8
73
74
71
69
70
72
43
44
67
68
45
46
65
66
63
7
64
59
60
61
6
62
5
2
19
20
17
18
1
11
12
30
32
29
31
41
42
13
14
10
4
5
54
58
57
53
56
55
52
51
60
6
59
